# Translation of the ERM-1 membrane-binding domain directs *erm-1* mRNA localization to the plasma membrane in the *C. elegans* embryo

**DOI:** 10.1101/2022.05.11.491546

**Authors:** Lindsay P. Winkenbach, Dylan M. Parker, Robert T. P. Williams, Erin Osborne Nishimura

## Abstract

mRNA localization and transport are integral in regulating gene expression. In *Caenorhabditis elegans* embryos, the maternally inherited mRNA *erm-1 (Ezrin/Radixin/Moesin)* concentrates in anterior blastomeres. *erm-1* mRNA localizes within those blastomeres to the plasma membrane where the essential ERM-1 protein, a membrane-actin linker, is also found. We demonstrate that the localization of *erm-1* mRNA to the plasma membrane is translation-dependent and requires its encoded N-terminal membrane-binding (FERM) domain. By perturbing translation through multiple methods, we found *erm-1* mRNA localization at the plasma membrane was maintained only if the nascent peptide remained in complex with the translating mRNA. Indeed, recoding the *erm-1* mRNA coding sequence while preserving the encoded amino acid sequence did not disrupt *erm-1* mRNA localization, corroborating that the information directing mRNA localization resides within its membrane-binding protein domain. A smiFISH screen of 17 genes encoding similar membrane-binding domains identified three plasma membrane localized mRNAs in the early embryo. Nine additional transcripts showed apparent membrane localization later in development. These findings point to a translation-dependent pathway for localization of mRNAs encoding membrane-associated proteins.

**SUMMARY STATEMENT:** In *C. elegans, erm-1* mRNA localization to plasma membranes requires translation of the ERM-1 membrane-binding domain

## INTRODUCTION

mRNA localization is a prevalent feature in diverse cell types and organisms (Chouaib et al., 2020; Knowles et al., 1996; Lécuyer et al., 2007; Llopis et al., 2010; Long et al., 1997; Rebagliati et al., 1985). Subcellular localization of mRNA is associated with spatiotemporal control of gene expression. mRNA localization can occur as a cause or consequence of translational regulatory control, can promote mRNA degradation, facilitate interactions with effector proteins, and prevent premature non-specific interactions (Besse and Ephrussi, 2008; Broadus et al., 1998; Chouaib et al., 2020; Ephrussi et al., 1991; Frohnhöfer and Nüsslein-Volhard, 1986; Ma and Mayr, 2018; Rebagliati et al., 1985; Ryder and Lerit, 2018; Sepulveda et al., 2018). In *Caenorhabditis elegans* early embryos, mRNA localization is a prominent feature of maternally inherited mRNAs and may contribute to cell-specific patterning prior to the onset of zygotic transcription (Aoki et al., 2021; Lee et al., 2020; Nishimura et al., 2015; Parker et al., 2020; Updike and Strome, 2010). Generally, maternal transcripts enriched in the posterior cells of the embryo localize to membraneless biomolecular condensates called P granules (Lee et al., 2020; Parker et al., 2020). Previous work has indicated that transcripts localize to P granules following translation repression (Lee et al., 2020; Parker et al., 2020). In contrast, maternal transcripts that concentrate in the anterior cells often localize to the plasma membrane (Parker et al., 2020) often colocalizing with their encoded proteins. However, the molecular mechanisms that facilitate membrane localization in *C. elegans* are unclear. Here we focus on *erm-1* as a model membrane-associated transcript and characterize the mechanisms underlying its localization to the cell membrane.

mRNA localization can occur via translation-independent or translation-dependent pathways (Johnston, 2005; Parton et al., 2014; Szostak and Gebauer, 2013). Translation-independent pathways typically rely on *cis-*acting elements, RNA sequences or structures that are often located in untranslated regions (UTRs) and recruit *trans-*recognition factors such as RNA Binding Proteins (RBPs). Recognition by RBPs can lead to either passive or directed transport, often in association with other processes such as mRNA protection, mRNA degradation, or translational regulatory control (Engel et al., 2020). The result is an enrichment of mRNAs in specific subcellular locales. In *C. elegans*, some posterior-enriched maternal transcripts localize through translation-independent pathways, relying on *cis*-acting elements in their 3’UTRs to direct translational repression that is required for localization into P granules (Lee et al., 2020; Parker et al., 2020). In contrast, translation-dependent pathways of mRNA localization typically rely on peptide signals in the nascent polypeptide. Transcripts that concentrate at the ER often rely on signal peptides to direct translating mRNAs and their encoded proteins to their destinations (Walter and Johnson, 1994). However, previously identified transcripts that localize to the plasma membrane in *C. elegans* embryos lack a discernable signal peptide (Parker et al., 2020). Recently, several *C. elegans* transcripts that encode members of the apical junction sub-complexes, their additional ancillary proteins, or other cytoskeletal components were found to localize to the plasma membrane during mid-embryogenesis with subcellular localization patterns that did not appear to overlap with the endoplasmic reticulum. Of these, the *dlg-1* transcript was shown to localize in a translation-dependent fashion (Tocchini et al., 2021). Together, these findings demonstrate the localization of plasma membrane enriched transcripts is occurring in a distinct manner from the canonical, ER signal peptide directed pathway and that local translation may be a general feature of junction and membrane-linker proteins.

*erm-1* (*ezrin/radixin/*moesin) mRNA is the most anterior-enriched transcript in the 2-cell *C. elegans* embryo (Nishimura et al., 2015; Tintori et al., 2016). In addition to its high enrichment within anterior cells of the early embryo, *erm-1* mRNA concentrates at plasma membranes within those cells, a pattern coincident with its encoded ERM-1 protein (Furden et al., 2004; Göbel et al., 2004; Parker et al., 2020). Previously, we showed the *erm-1* 3’UTR was insufficient to direct membrane mRNA localization, indicating that the localization element resides elsewhere in the RNA sequence or the encoded protein (Parker et al., 2020). In this study, we set out to identify which elements in *erm-1* mRNA or the encoded ERM-1 protein are necessary for membrane localization.

In *C. elegans*, ERM-1 is the sole ortholog of the conserved ERM protein family that serve as membrane-actin linkers (Furden et al., 2004). ERM proteins regulate cell morphology and signaling events at the plasma membrane. Therefore, they are prominent in processes such as epithelial junction remodeling, cell migration, promotion of microvilli formation, and interactions with actin at the cell cortex (Fehon et al., 2010; Furden et al., 2004; Göbel et al., 2004; McClatchey, 2014). Proper specialization of the cell cortex and plasma membrane is critical for controlling cell morphology, as evidenced by the fact that in *C. elegans*, loss of the *erm-1* in the intestine results in early embryo lethality due to constrictions and disjunctions in the intestinal lumen (Furden et al., 2004; Ramalho et al., 2020).

Here we demonstrate *erm-1* mRNA accumulation at the plasma membrane is translation-dependent and requires the membrane-binding ability of the FERM domain to enrich at the plasma membrane. Further, we screened 17 genes encoding similar membrane-binding FERM or PH-like domains. We identified twelve additional plasma membrane localized transcripts and other patterns of subcellular mRNA localization that change over developmental time. Our findings suggest translation of this conserved membrane binding domain is conducive to subcellular localization of both the mRNA and the encoded protein.

## RESULTS

### *erm-1* mRNA localization to the plasma membrane requires translation initiation

mRNA localization is directed through either translation-dependent or translation-independent pathways (Johnston, 2005; Parton et al., 2014; Szostak and Gebauer, 2013). To test which pathway was responsible for *erm-1* mRNA localization, we disrupted global translation by two methods and determined whether either perturbed *erm-1* mRNA accumulation at the membrane. We first depleted the translation initiation factor *ifg-1 (Initiation Factor 4G (eIF4G) family)* by RNA interference (RNAi). IFG-1 is the sole *C. elegans* ortholog of eIF4G, and both cap-dependent and -independent translation initiation require IFG-1 (Berset et al., 1998; Kim et al., 2018; Ramírez-Valle et al., 2008; Rogers et al., 2011). Using a destabilized-GFP as a translation reporter (MODCPEST GFP::H2B), we found that *ifg-1* RNAi decreased translation in a partially penetrant fashion as indicated by a decrease in GFP::H2B fluorescence (Corish and Tyler-Smith, 1999; Kaymak et al., 2016; Li et al., 1998) (**Fig. 1A, C**). *ifg-1* RNAi introduced in the L2 stage of development led to 46% of 4-cell progeny exhibiting a significant loss of GFP signal and 54% showing no significant change compared to wild type. Importantly, we observed that embryos with impacted translation also experienced a loss of *erm-1* mRNA localization at the plasma membrane with high concordance (**Fig. 1A, C**). These results support the model that *erm-1* mRNA localization to the plasma membrane is translation-dependent.

**Figure 1:**
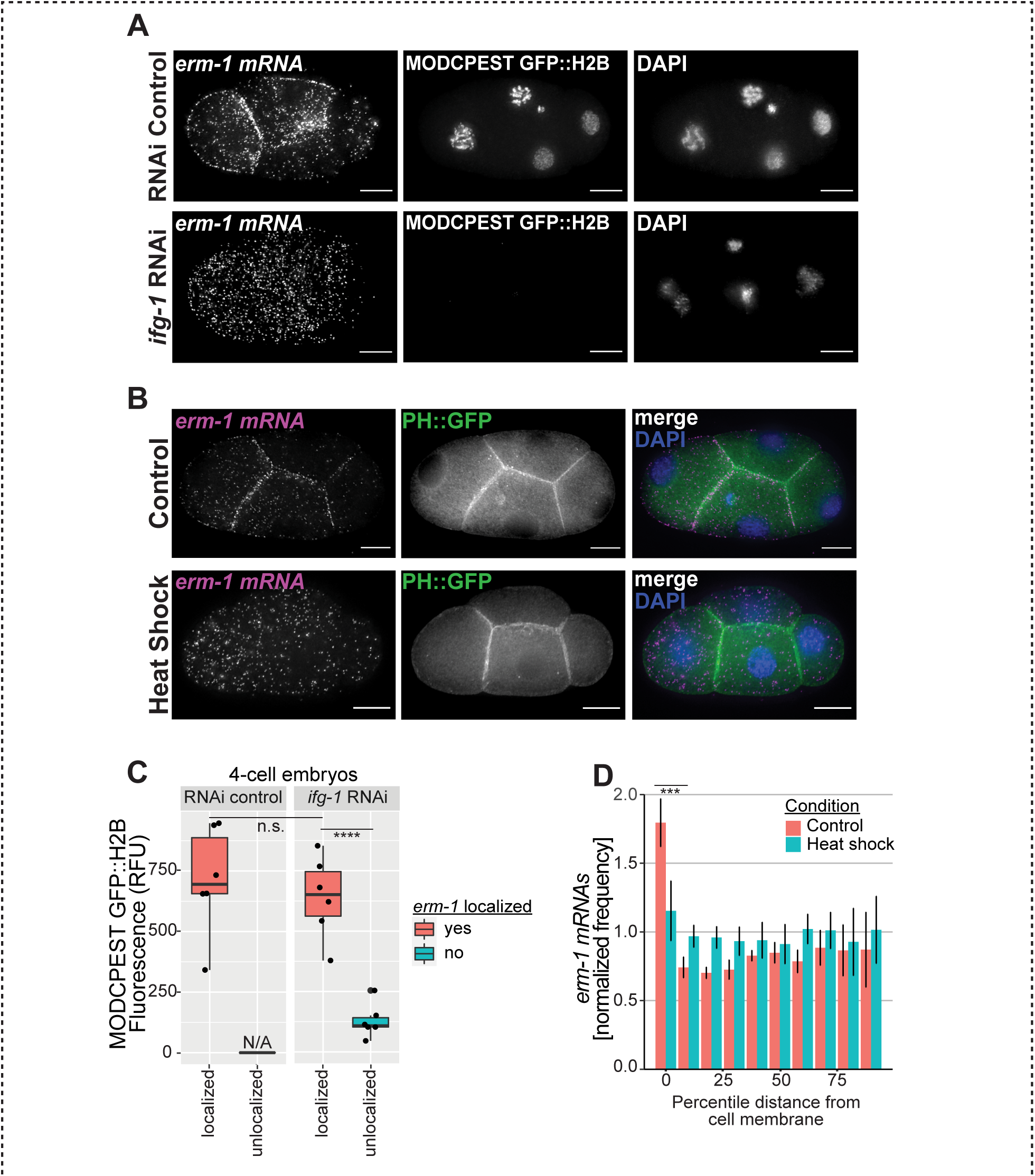
Disruption of global translation leads to loss of *erm-1* mRNA localized to the membrane. **(A)** Fluorescent micrographs of 4-cell stage *C. elegans* embryos. RNAi was performed to deplete the translational initiation factor *ifg-1*, or an empty vector RNAi control was performed. *erm-1* transcripts were imaged by smFISH. In the same embryos, GFP from a translational reporter transgene (MODCPEST-GFP::H2B) and DNA (DAPI) were also imaged. Representative images are shown from a total of 102 4-cell stage embryos surveyed (*n = 23* RNAi control, *n = 79 ifg-1* RNAi [*n = 43* reporter signal retained, *n = 36* reporter signal depleted]). Scale bars are 10 μm. **(B)** 4-cell stage *C. elegans* embryos harboring a membrane marker transgene (PH::GFP, green) were imaged for *erm-1* transcripts by smFISH (magenta) under no heat shock (control 20°C) and heat stress conditions (25 minutes at 30°C heat shock). DNA was also imaged as DAPI staining (blue). Scale bars are 10 μm. **(C)** Quantification of translation reporter fluorescence under RNAi control (*n = 6*), *ifg-1* RNAi *erm-1* localization retained (*n = 6*), *ifg-1* RNAi *erm-1* localization lost (*n = 6*) conditions. Background subtracted GFP intensities were measured as relative fluorescent units (RFU) using nuclear masks generated using DAPI. *erm-1* mRNA localization was assessed qualitatively as localized or unlocalized in 4-cell embryos. Significance indicates *P*-values derived from Welch Two Sample *t*-tests comparing the RFU values for localized versus unlocalized for the transcript *erm-1* at the given condition. P value legend: 0.00005>\*\*\*\***(D)** Quantification of *erm-1* mRNA under control (a representative set of *n=5*) and heat shock conditions (a representative set of *n=7*) indicating the normalized frequency of *erm-1* mRNA at increasing, normalized distances from the cell periphery. Significance indicates *P*-values derived from Welch Two Sample *t*-tests comparing the cell membrane localization for heat shock versus control for the transcript *erm-1* at the given stage. *P* value legend: ***>0.00005

As a complementary approach to disrupting global translation initiation via RNAi, we next disrupted translation through heat shock and quantified the resulting changes in *erm-1* mRNA enrichment at the plasma membrane (Parker et al., 2020). Heat shock prevents protein synthesis primarily through changes in phosphorylation states of translation initiation factors followed by their subsequent inactivation (Cuesta et al., 2000; Duncan and Hershey, 1984; Shalgi et al., 2013). Heat shock acts within a shorter time frame than *ifg-1* RNAi (25 minutes heat shock vs. 48 hrs RNAi exposure). In heat treated 4-cell embryos, we observed a 1.6-fold reduction in *erm-1* mRNA enrichment alongside the plasma membrane (at a distance of less than 10% of the normalized radius from the plasma membrane) after only 25 minutes of 30 °C heat exposure compared to controls that were kept at 20 °C for the same duration (**Fig. 1B, D**). Combined with the *ifg-1* RNAi experiment findings, this illustrated that *erm-1* mRNA localization to the plasma membrane depended on translation-initiation for establishment and maintenance. Together, these results suggest a translation-dependent pathway and imply that the signal to localize *erm-1* mRNA to membranes may be an encoded peptide sequence in the nascent ERM-1 protein. However, these assays do not yield information on whether active translation or an intact ribosome nascent chain complex (RNC), or both, are required for localization.

### The ERM-1 nascent peptide is required for *erm-1 mRNA* enrichment at plasma membranes

We identified that translation is required for *erm-1* mRNA enrichment at plasma membranes. This suggests a model in which the RNC – comprised of *erm-1* mRNA, the translating ribosome, and the emerging nascent ERM-1 protein – is transported to the plasma membrane together through recognition of amino acid sequences in the nascent ERM-1 protein. We hypothesize that *erm-1* mRNA localization requires intact RNCs, likely at steps that both establish and maintain localization. To test this hypothesis, we inhibited translation elongation using two different drugs, one that preserves RNCs (cycloheximide) and one that disrupts them (puromycin) (Azzam and Algranati, 1973; Schneider-Poetsch et al., 2010).

The eggshell and permeability barrier in the *C. elegans* embryo complicate drug treatment by limiting small molecule penetrance (Carvalho et al., 2011; Olson et al., 2012; Stein and Golden, 2015). To circumvent this, we disrupted the sugar modifying enzyme and permeability barrier protein PERM-1 by RNAi, thereby allowing ingression of small molecules such as cycloheximide and puromycin. Though *perm-1* RNAi eventually leads to lethality in late embryos, development in early embryonic stages proceeds typically (Carvalho et al., 2011). Importantly, *perm-1* RNAi is compatible with both drug treatment and smFISH imaging of *erm-1* mRNA.

We observed that disruption of the RNC via puromycin treatment led to loss of *erm-1* mRNA localization at the membrane in 84% of embryos between the 2-cell and 8-cell stages (**Fig. 2A, B; Fig S1A, B**). In contrast, cycloheximide treatment, which stalls translation during elongation while preserving the RNC only altered *erm-1* mRNA localization in 4% of embryos surveyed (**Fig. 2A, B; Fig S1A, B**). This suggests that the *erm-1* mRNA must maintain association with the ribosome for *erm-1* mRNA molecules to maintain localization to plasma membranes upon translation disruption. Additionally, the maintenance of *erm-1* mRNA localization does not require ongoing translational elongation provided stalled RNCs are preserved intact, as is the case with cycloheximide treatment. These findings further support the translation-dependent model and suggest that *erm-1* mRNA transcripts localize through association with the RNC.

**Figure 2:**
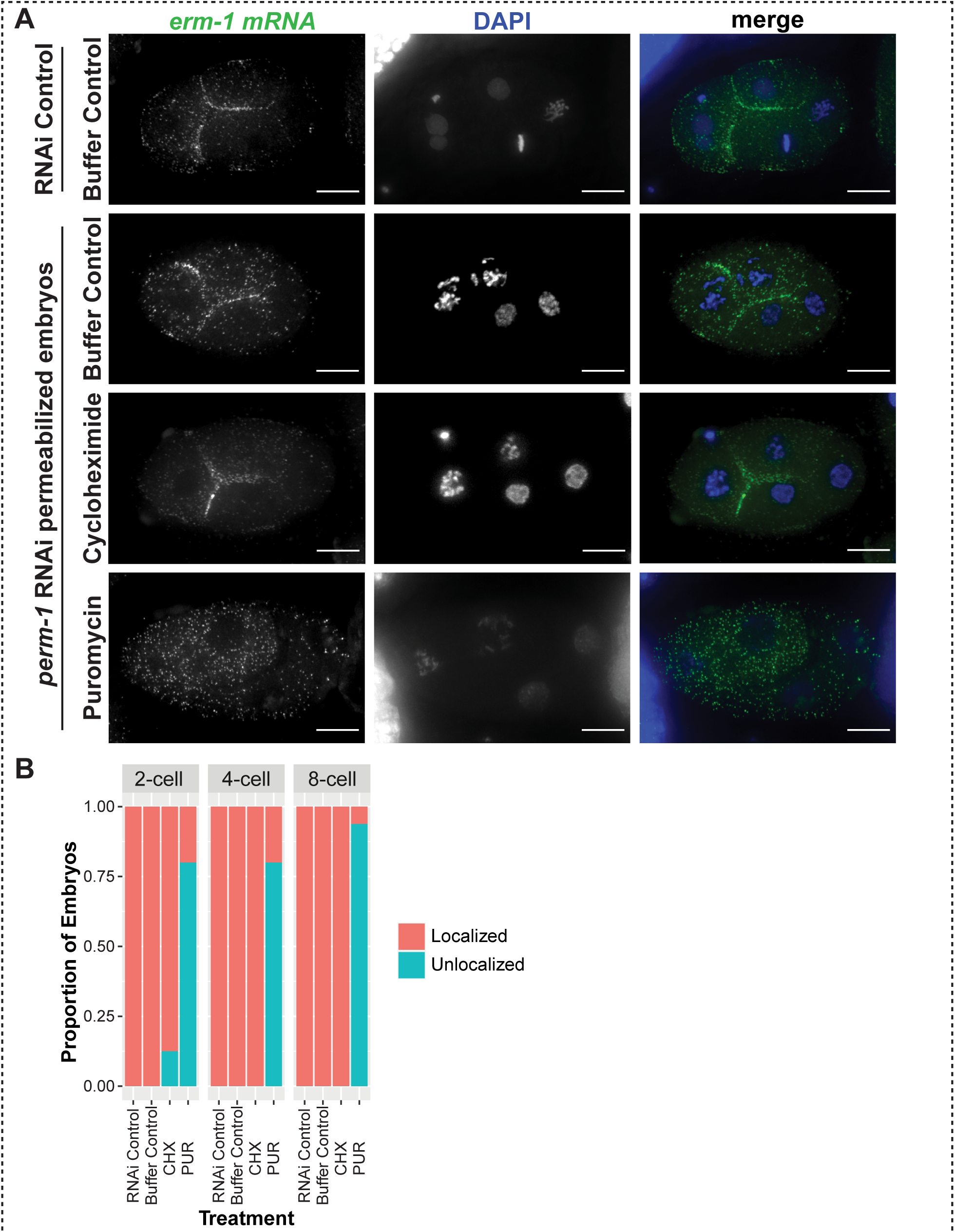
The intact ERM-1 ribosome nascent chain complex (RNC) is required for *erm-1* mRNA enrichment at cell membranes. **(A)** Fluorescent micrographs of 4-cell stage *C. elegans* embryos are shown in which embryos were permeabilized by *perm-1* RNAi and subsequently treated with small molecule translation inhibitors, in comparison to RNAi and drug treatment controls. *erm-1* mRNA (green) was imaged by smFISH, under control, cycloheximide (500 μg/mL, 20min), or puromycin (500 μg/mL, 20min) treatment conditions. Scale bars 10 μm. **(B)** Bar plot indicating the proportion of embryos displaying *erm-1* mRNA enriched at cell membranes (localized) or homogenously distributed through the cell (unlocalized) for 2-cell, 4-cell, and 8-cell embryos subjected to the indicated treatments.

### *erm-1* mRNA localization to the plasma membrane does not depend on nucleic acid sequences

We have established that *erm-1* mRNA localizes to plasma membranes in a translation-dependent manner. However, *erm-1* mRNA can persist at membranes if the RNC remains intact when disrupting translation elongation. This suggests that localization is dependent on ERM-1 amino-acid sequence and not *erm-1* mRNA sequence. Supporting this hypothesis, previous evidence has illustrated that the *erm-1* 3’UTR (typically a common site of *cis*-acting localization elements) is insufficient to direct mRNA to the membrane (Parker et al., 2020). To test whether other *erm-1* mRNA nucleotide sequences are dispensable for localization, we artificially recoded the *erm-1* mRNA nucleotide sequence while preserving the amino acid sequence by capitalizing on the redundancy of the genetic code (**Fig S2A)**. Our nucleotide recoded, yet amino-acid synonymous, *erm-1* sequence (called *erm-1 synon*) shares 64% identity at the nucleic acid level with the wild-type *erm-1* sequence (called *erm-1*) while maintaining 100% identity of the amino acid sequence (**Fig. 3A; Fig S2B**. We designed single molecule inexpensive FISH (smiFISH) probes that could distinguish between the recoded, synonymous *erm-1* and wild-type sequences (**Fig. 3B, C)**. Using these probes, we found the *erm-1 synon* transcript retained enrichment at the plasma membrane with no significant difference between it and either the endogenous *erm-1* transcript (**Fig. 3B D**) or a matched transgenic wild-type *erm-1* sequence inserted at the same transgenic location (**Fig. S2D, C)**. These data imply that RNA sequences within the *erm-1* transcript are dispensable for *erm-1* mRNA localization, and instead, localization elements reside in the translated ERM-1 protein.

**Figure 3:**
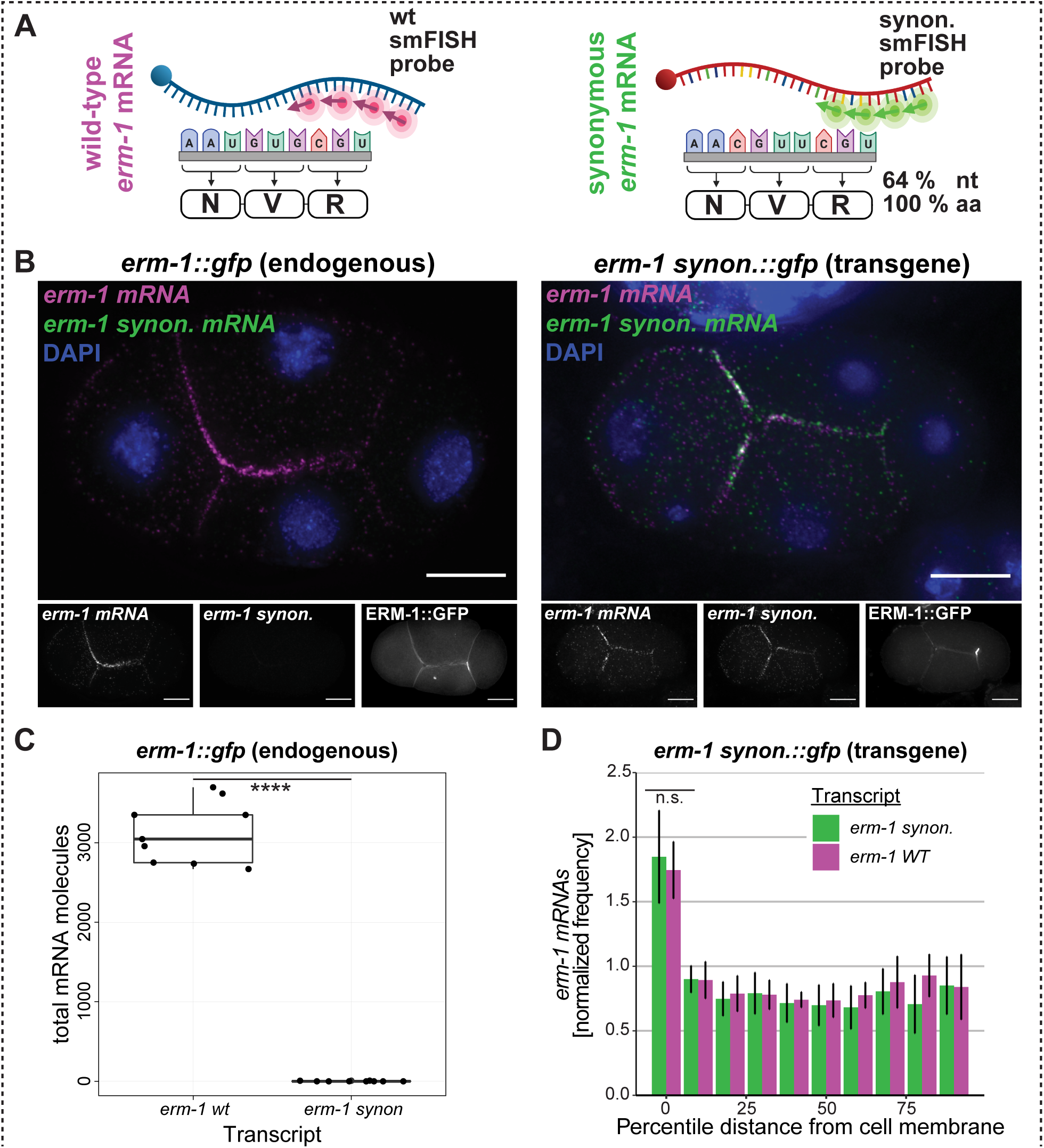
*erm-1* mRNA localization to the cell membrane is mRNA coding sequence independent. **(A)** Schematic depictions comparing wild-type *erm-1* mRNA (magenta) to the recoded, *synonymous erm-1* mRNA (green). **(B)** smFISH micrographs of 4-cell staged *C. elegans* embryos imaging wild-type *erm-1* (magenta) and the re-coded, *synonymous erm-1* mRNA (*erm-1 synon. mRNA*, green). GFP tagged *erm-1* control 4-cell embryo (left) and MosSCI GFP tagged *synonymous erm-1* 4-cell embryo (right). DNA (DAPI, blue) and membranes marked by ERM-1::GFP are also shown. Scale bars 10 μm. **(C)** Total number of *erm-1 wt* and *erm-1* synonymous mRNA molecules detected in *erm-1*::*gfp* (endogenous) (*n = 9*). Significance indicates *P*-values derived from Welch Two Sample *t*-tests comparing the total number of *erm-1* molecules detected by the *erm-1 wt* probe vs the *erm-1 synonymous* probe in the endogenously tagged *erm-1::gfp* background. **(D)** Quantification of endogenous and synonymous *erm-1* mRNA (*n = 6*) indicating the normalized frequency of mRNAs within binned, percentile distances from the cell membrane counted and normalized against the total volume of each cell. Significance indicates *P*-values derived from Welch Two Sample *t*-tests comparing the cell membrane localization of endogenous and synonymous *erm-1* mRNA in the *erm-1 synon::gfp* transgenic background. *P* value legend: NS>0.05; ***>0.00005

### *erm-1* mRNA and ERM-1 protein localization require the ERM-1 PIP_2_ membrane-binding region

To identify domains within the ERM-1 protein required to localize translating *erm-1* to plasma membranes, we examined *erm-1* mRNA localization upon mutations in key conserved ERM-1 domains. Generally, ERM proteins are a conserved family defined by domains common to their founding members, Ezrin, Radixin, and Moesin (Bretscher, 1983; Lankes and Furthmayr, 1991; Tsukita et al., 1989). ERM proteins serve as structural linkers between the plasma membrane and the actin cytoskeleton and play central roles in cell morphology and signaling processes that converge on the plasma membrane. Two key domains coordinate their linker function. The N-terminal band 4.1 Ezrin/Radixin/Moesin (FERM) domain houses a PH-like (Pleckstrin Homology-like) domain that associates with the plasma membrane through interactions with PIP_2_ (phosphatidylinositol (4,5) bisphosphate) (Barret et al., 2000; Fehon et al., 2010; Roch et al., 2010). In contrast, the C-terminal C-ERMAD domain interacts with the actin cytoskeleton in a phosphorylation-dependent manner (McClatchey, 2014; Ramalho et al., 2020). The FERM and C-ERMAD domains can also intramolecularly bind to prevent their respective substrate associations. A dephosphorylation event on the C-ERMAD domain increases intramolecular affinity thereby permitting the inactive form (Li et al., 2007; Pearson et al., 2000; Roch et al., 2010). Therefore, the architecture of ERM-1 connects the plasma membrane and the actin cytoskeleton in a phosphorylation-dependent mechanism.

Mutating four lysines to asparagines abrogates the PIP_2_ binding ability of the FERM domain (Barret et al., 2000; Roch et al., 2010) (**Fig. 4A**), termed the ERM-1[4KN] mutant (Ramalho et al., 2020). In *C. elegans*, this leads to intestinal lumen cysts and disjunctions as well as early larval lethality that phenocopy the *erm-1* null (Göbel et al., 2004; Ramalho et al., 2020). In contrast, mutating the conserved, phosphorylatable residue T544 to alanine (ERM-1[T544A]) disrupts the function of the C-ERMAD domain, thereby rendering C-ERMAD non-phosphorylatable (Carreno et al., 2008; Zhang et al., 2020). In *C. elegans*, this leads to disrupted cortical actin organization and reduced apical localization of ERM-1 (Ramalho et al., 2020). We assessed *erm-1* mRNA localization in these two previously characterized mutant strains to determine whether *erm-1* mRNA accumulation at the plasma membrane requires the FERM or C-ERMAD domains.

**Figure 4:**
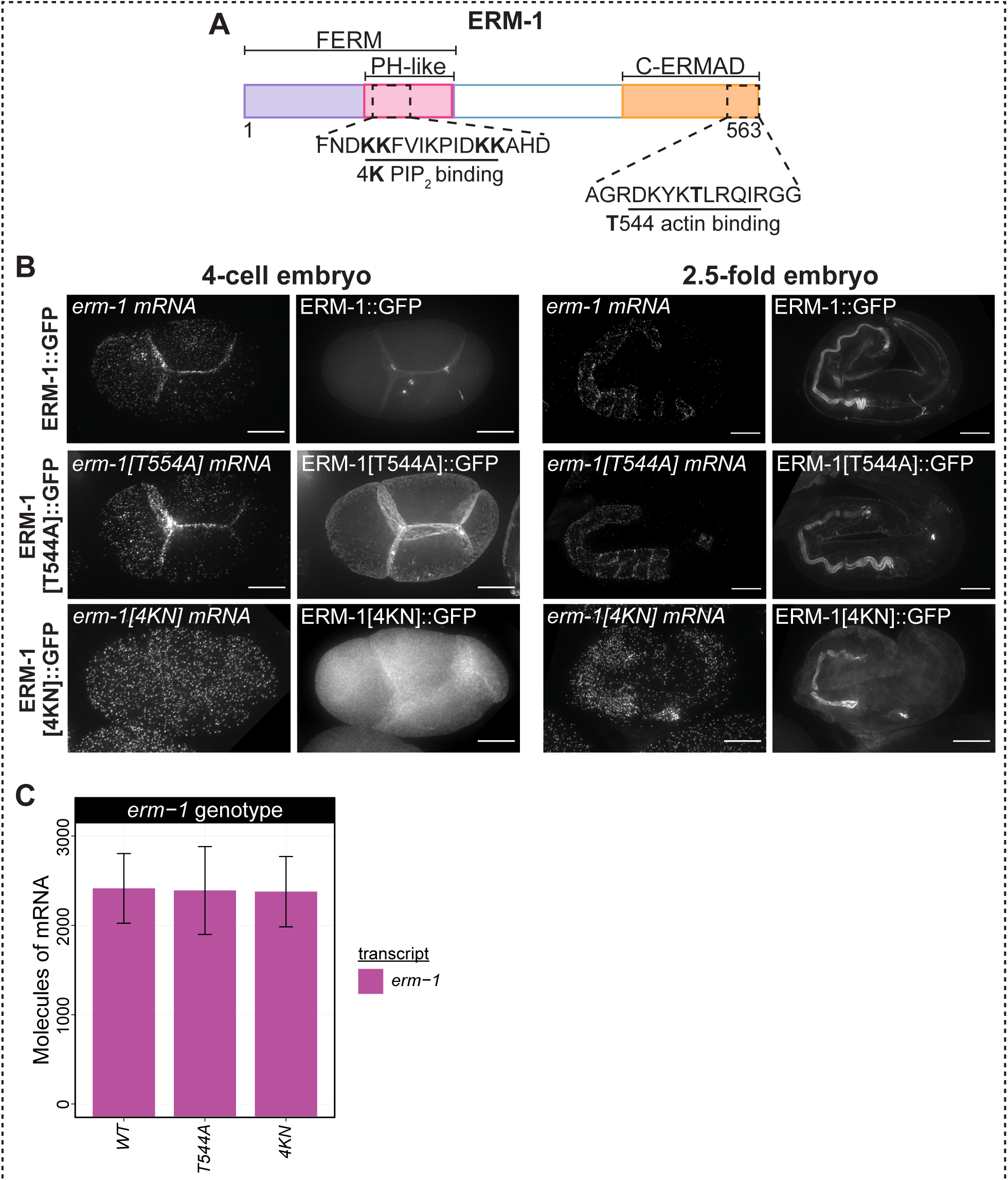
PIP_2_ binding is required for *erm-1* mRNA and ERM-1 protein localization in early and late embryos. **(A)** Schematic model of the ERM-1 protein showing the N-terminal FERM domain (purple) containing the conserved PIP_2_ – membrane binding region (PH-like, pink) and C-terminal conserved C-ERMAD actin-binding domain (orange). **(B)** smFISH micrographs of 4-cell and 2.5-fold embryos displaying *erm-1* mRNA and ERM-1 protein localization in *erm-1::gfp, erm-1[T544A]::gfp*, and *erm-1[4KN]::GFP* backgrounds. Scale bars 10 μm. **(C)** *erm-1* mRNA abundance does not significantly differ between *erm-1::gfp, erm-1[T544A]::gfp*, and *erm-1[4KN]::GFP* expressing strains. *P*-values derived from Welch Two Sample *t*-tests comparing the number of *erm-1* mRNA molecules detected in the *erm-1::gfp, erm-1[T544A]::gfp*, and *erm-1[4KN]::GFP* expressing strains. *P* value legend: NS>0.05.

Because the ERM-1[T544A] and ERM[4KN] mutant strains were extensively studied in previous reports during mid-stage embryogenesis for their impacts on intestinal development (Ramalho et al., 2020), we examined *erm-1* mRNA localization at both the 4-cell stage we have previously studied and at mid-stage embryogenesis. These assays were performed in ERM-1[T544A] homozygous mutants with no wild-type ERM-1 in the background. In 4-cell stage embryos, both *erm-1[T544A]* mRNA and ERM-1[T544A] protein localized to the plasma membranes similar to wild-type (**Fig. 4B)**. However, the mutants exhibited plasma membranes with “ruffled” or distorted phenotypes, indicating that loss of this phospho-moiety imparts a cellular phenotype through its previously reported reduction in actin organization (**Fig. 4B)** (Ramalho et al., 2020). In 2.5-fold embryos, *erm-1[T544A]* mRNA also localized at the plasma membrane, similar to wild-type strains. In these embryos, the ERM-1[T544A] protein displayed increased basolateral localization in the intestinal lumen, as has been previously reported (**Fig. 4B)** (Ramalho et al., 2020). Therefore, the T544A mutant, though disruptive of ERM-1 phosphorylation and actin association, was not sufficient to disrupt *erm-1* mRNA localization to the plasma membrane at these stages.

Mutations in ERM-1 that disrupt FERM domain/PIP_2_ interactions (called ERM-1[4KN]) reduce (in the intestine and excretory canal) or eliminate (in progenitor germ cells and seam cells) ERM-1[4KN] protein localization to apical membranes (Ramalho et al., 2020). We performed smFISH on 4-cell and 2.5-fold stage ERM-1[4KN] homozygous mutant embryos lacking any ERM-1 wild-type copies to assess how this mutation impacted *erm-1* mRNA localization (**Fig. 4A; Fig S3A, B)**. At the 4-cell stage, *erm-1[4KN]* mRNA failed to localize to the plasma membrane in 100% of embryos surveyed **(Fig. 4B)**. However, the number of *erm-1* mRNA molecules detected did not significantly change between *erm-1 wt* and *erm-1[4KN]* (**Fig. 4C**). ERM-1[4KN] protein enrichment at the plasma membrane was also abolished. RNA localization was not observed at the 2.5-fold stage and protein localization was reduced, displaying disjunctions in the lumen as previously reported (Ramalho et al., 2020). Thus, the FERM-domain was required to localize both the *erm-1* mRNA and the ERM-1 protein to the plasma membrane. Combined with our previous findings, this indicates that the peptide signal required to localize the ERM-1 RNC, including its associated mRNA, resides within the FERM domain of the nascent peptide.

### Genes encoding FERM and PH-like domains are conducive for mRNAs with localization at the plasma membrane

Given that PIP_2_-binding FERM domains have a high affinity for membranes (Senju et al., 2017), their ability to direct membrane-localized mRNA transcripts may be generalizable. Evidence for this exists in other systems: In early *Drosophila* embryogenesis, the PIP_2_-binding PH domain and actin-binding domains of the Anillin protein are required to localize *anillin* mRNA to pseudocleavage furrow membranes (Hirashima et al., 2018). Based on the findings from the ERM-1[4KN]::GFP mutant strain, we hypothesized that the PIP_2_-binding element of the FERM domain could be a general predictor of transcripts that enrich at cell membranes.

To test our hypothesis, we conducted a smiFISH-based screen in *C. elegans* early embryos for membrane localization of other PIP_2_-binding FERM or PH-like domain containing transcripts (Nishimura et al., 2015; Tintori et al., 2016). A total of 17 transcripts (9 with FERM domains and 8 with PH-like domains) were selected for visualization based on their expression in early embryos and with preference given to homology in other organisms **(Table 1; Fig S6; Fig S7; Materials and Methods)**. The set of transcripts selected comprised those that enriched to anterior embryonic cells, posterior embryonic cells, and those with ubiquitous distribution across the early embryo.

Of the 17 screened transcripts, three (*frm-4, frm-7, ani-1)* displayed clear membrane localization in early embryos at the 4-cell stage (**Fig. 5A, B; Fig. S4A-C)**. Seven transcripts displayed clustered subcellular patterning in the posterior cell (*Y41E3*.*7, unc-112, mrck-1, wsp-1, ani-2, dyn-1*, and *exoc-8*) (**Fig. S4C**). Four transcripts displayed uniform distribution or other patterns (*F07C6*.*4, efa-6, mtm-1*, and *let-502*) (**Fig. S4A**). Three transcripts (*frm-2, nfm-1*, and *ptp-1*) yielded RNA abundance too low to determine subcellular enrichment (**Table 1; Fig. S4B**).

**Figure 5:**
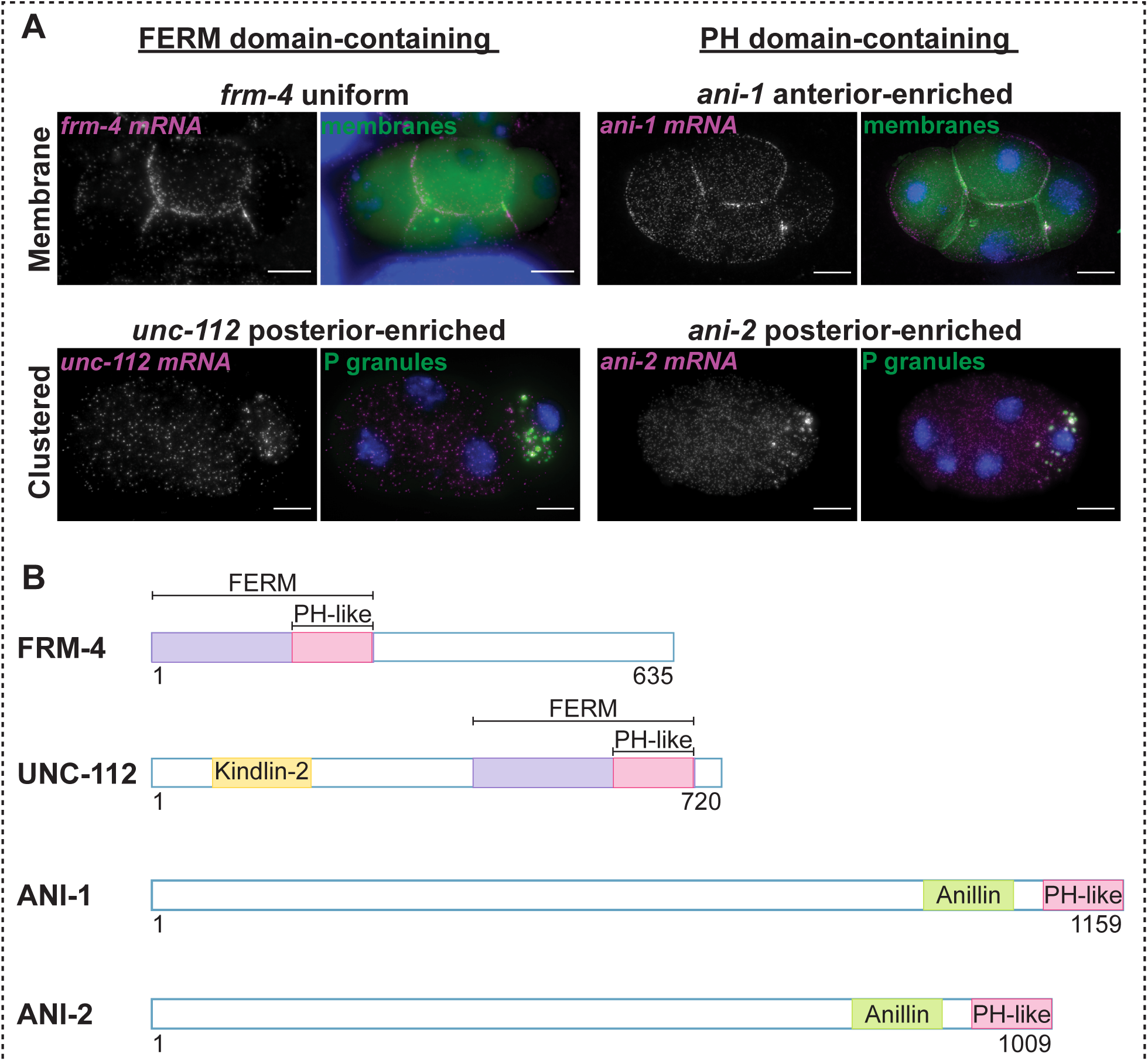
FERM and PH domain-containing, maternally-inherited transcripts display mRNA patterning. **(A)** smFISH micrographs of 4-cell embryos imaging two FERM domain containing transcripts, *frm-4, unc-112*, (magenta, left) and two PH-domain containing transcripts *ani-1, ani-2* (magenta, right). *frm-4* mRNA and *ani-1* mRNA (top) were imaged in a PH::GFP membrane marker transgenic background (membranes, green). *unc-112* mRNA and *ani-12* mRNA (bottom) were imaged in the GLH-1::GFP P granule marker strain (P granules, green). DNA (DAPI, blue). Scale bars 10 μm.

Notably, all transcripts previously reported as posterior-cell enriched (Nishimura et al., 2015; Tintori et al., 2016) exhibited clustered patterning in early embryos (**Table 1; Fig. S4C**). This is consistent with prior visualizations of posterior-enriched transcripts at the 4-cell stage that often concentrate within P granules (Parker et al., 2020). Of the transcripts that were uniformly distributed between the anterior and posterior cells (as assayed by RNA-seq), those with posterior-enrichment below the significance cut-off (*Y43E3*.*7, mrck-1, wsp-1*) also displayed clustered patterning (**Table 1; Fig. S4C**). Overall, these results support previous findings in the early embryo that AB-cell enriched transcripts tend to localize to membranes, whereas P_1_-enriched transcripts tend to localize to P granules, membraneless organelles housing lowly translated mRNAs (Lee et al., 2020; Parker et al., 2020; Updike and Strome, 2010). To determine whether clustered transcripts were indeed localizing to P granules or membranes, smFISH was conducted in a P granule or membrane marker strain, respectively. Indeed, even in the case of the homologs *ani-1* and *ani-2*, the anterior-enriched *ani-1* mRNA was membrane localized, whereas the posterior-enriched *ani-2* mRNA localized to P granules (**Fig. 5A**). Importantly, ANI-1 protein is translated in early embryos and concentrates at the plasma membrane while early embryo expression of ANI-2 protein is not detected (Maddox et al., 2005). Together, these results suggest that translational dependence of plasma membrane mRNA localization is not limited to *erm-1*.

Our visual screen found that subcellular localization patterns of some transcripts changed over developmental time. At the 4-cell stage, *unc-112* mRNA clustered in the P-lineage in P granules (**Fig. 5A**). However, later in embryonic development, at the 3-fold stage, *unc-112* mRNA localized along the body wall where the encoded UNC-112 protein reportedly functions (**Fig. S5A**) (Rogalski et al., 2000). At the 4-cell stage, *F07C6*.*4* mRNA had uniform distribution (**Fig. 5A**), but at the 100-cell stage it localized to the plasma membranes of a subset of discrete cells (**Fig. S5B**). Similarly, at the 4-cell stage *let-502* had uniform localization (**Fig. S4B**), but at the 1.5-fold stage it was localized to adherens junctions where the encoded LET-502 protein localizes and functions (**Fig. S5C**) (Piekny et al., 2003). While the *ani-2* transcript clustered in P granules at the 4-cell stage (**Fig. 5A**), at the bean stage *ani-2* was excluded from P granules and found at the interface of the primordial germ cells where ANI-2 functions in maintaining the structure of the syncytial compartment of germline cytoplasm at the membrane (Maddox et al., 2005) (**Fig. S5D**). Overall, 9 of the surveyed transcripts without observable membrane localization at the 4-cell stage (*F07C6*.*4, frm-2, nfm-1, ptp-1, unc-112, efa-6, mtm-1, let-502, ani-2*, and *dyn-1*) displayed cortical localization at later stages of development, frequently coinciding with where their encoded proteins function (Hunt-Newbury et al., 2007; Josephson et al., 2017; Maddox et al., 2005; Piekny et al., 2003; Rogalski et al., 2000; Thompson et al., 2002; Uchida et al., 2002; Zou et al., 2009). These findings suggest subcellular patterning is developmentally dynamic.

By conducting a visual screen on a subset of FERM and PH-like domain containing transcripts, we identified additional transcripts that enrich at membranes, adding to the small but growing list of transcripts with this behavior in *C. elegans* (Parker et al., 2020; Tocchini et al., 2021). 12 out of the 17 (70%) FERM and PH-like domain containing transcripts we surveyed exhibited membrane localization at some stage during development. These findings illustrate that translated mRNAs encoding PIP_2_-binding domains often localize to plasma membranes and suggests that they may locally translated at the sites where their proteins are required. Furthermore, results from this screen reinforce the trend that transcripts concentrated within the P lineage in the early embryo are lowly translated and display clustered patterning, likely within P granules.

## DISCUSSION

Here, we report that *C. elegans erm-1* mRNA is localized to the plasma membrane in a translation-dependent manner during early embryonic development (**Fig. 6**). We showed that in the absence of active translation, an intact ribosome-nascent chain complex (RNC) was required to maintain *erm-1* mRNA localization. We also demonstrated that 36% of the *erm-1* mRNA sequence could be altered but the transcript would still localize properly provided the ERM-1 protein sequence was preserved. This finding further suggests that the localization determinant is specified in the nascent chain of the ERM-1 protein, not as a *cis*-acting element in the mRNA sequence or structure. Furthermore, we narrowed down a domain required for localization and determined it resided within the FERM domain and depended on that domain’s ability to bind PIP_2_. We identified 3 additional FERM or PH-like domain encoding genes (*frm-7, frm-4*, and *ani-1*) with mRNA localization at the plasma membrane in the early embryo by a smiFISH visual screen and an additional 9 genes with mRNA localization later in development. Our data indicate subcellular localization is a generalizable feature of transcripts encoding FERM or PH-like domains.

**Figure 6:**
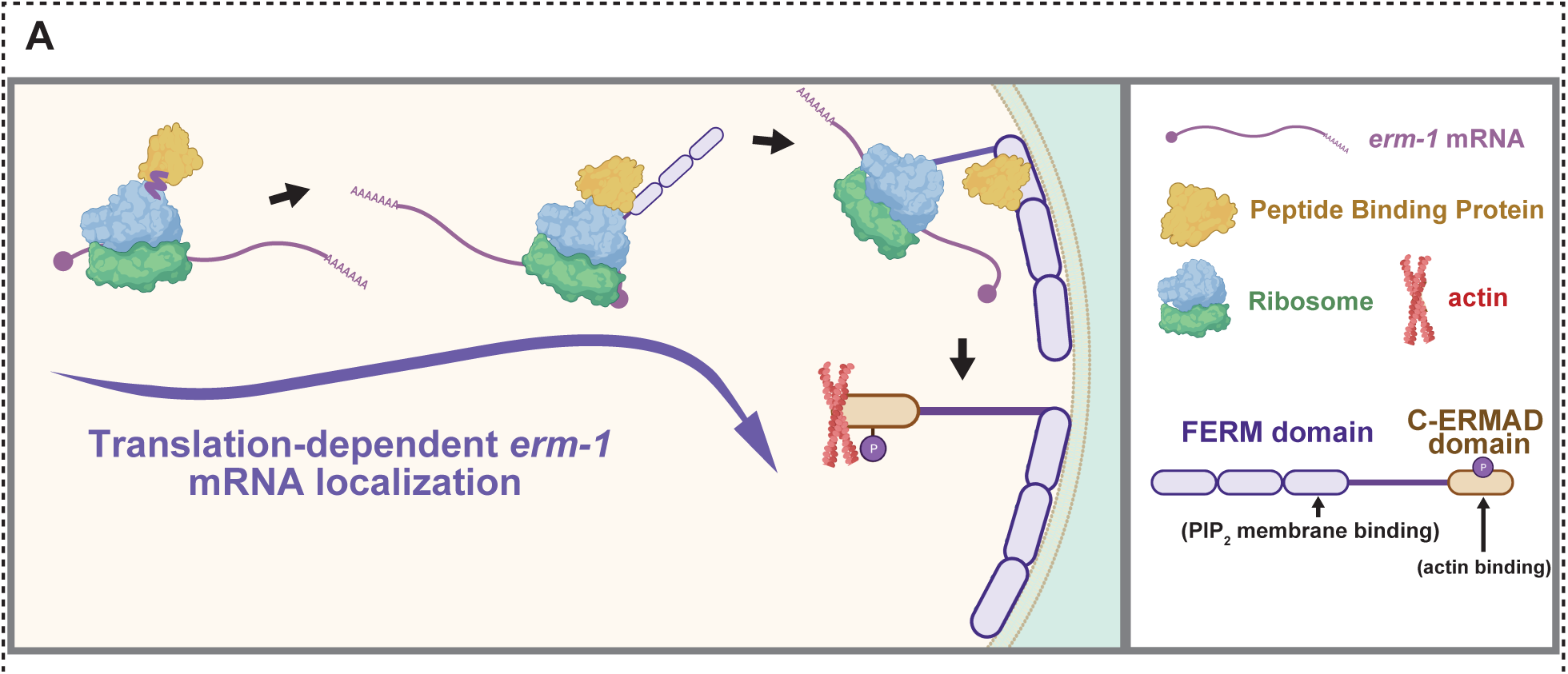
Model of translation-dependent *erm-1* mRNA localization in the early *C. elegans* embryo. *erm-1* mRNA is localized in a translation-dependent manner requiring the intact RNC and PIP_2_-binding region of the FERM domain.

Complementarily to our findings in the early embryos, recent studies of later developmental stages identified 8 transcripts (*dlg-1, ajm-1, sma-1, vab-10a, erm-1, pgp-1, magu-2*, and *let-413*), including *erm-1*, that enrich to regions of the plasma membrane adjacent to apical junctions (Li et al., 2021; Tocchini et al., 2021). In particular, *dlg-1* mRNA localizes through a translation-dependent pathway (Tocchini et al., 2021). *dlg-1* localization requires the translation of N-terminal L27-PDZ domains, C-terminal SH3, Hook, and Guk domains to fully recapitulate the localization patterns of the *dlg-1* mRNA. These data, in combination with our findings, suggest translation-dependent localization could be a prevalent feature of mRNA that generally encode apical junction components or membrane associated proteins.

An outstanding question is whether *erm-1* mRNA localization is critical for ERM-1 function. The ERM-1[4KN] mutation yields *erm-1* mRNA and ERM-1 protein that are mis-localized and results in lethality. However, the localization of the mRNA and protein are confounded, and it is difficult to test whether the proper localization of *erm-1* mRNA is required. Do ERM-1 proteins need to be locally translated at the plasma membrane for proper function? Because ERM proteins link the plasma membrane and actin cytoskeleton, ERM proteins often function in cell movement, membrane trafficking, cell signaling, and cell adhesion. They contribute to cancer-associated processes such as cell metastasis and chemotherapy resistance (Fehon et al., 2010; Kobori et al., 2021; McClatchey, 2003). Local translation of ERM-1 could be important for producing ERM-1 linker proteins at the exact sites and at the exact concentrations in which they are needed. This process could be responsive to signaling, polarity, or stability cues. Indeed, during embryonic development, the landscape of the plasma membrane is constantly changing and coordination between cell membrane and actin cytoskeletal structures are of paramount importance to cell morphology and cell migration processes. Therefore, local translation of ERM-1 could be sensitive to incoming signals or developmental needs.

As an example of this concept, the protein PCNT (Pericentrin) in human cells and zebrafish embryos is shown to cotranslationally localize to dividing centrosomes during early mitosis (Sepulveda et al., 2018). This is hypothesized to supply sufficient PCNT protein where it is expeditiously needed for cell division and to mitigate the kinetic challenge of trafficking a large protein. Alternatively, some proteins are locally translated to promote proper folding, facilitate interactions with effector proteins, or to promote their ultimate integration into membranes or vesicles (Das et al., 2021). We postulate another idea, that local translation may temporarily stabilize ERM associations at membranes until phosphorylation in the C-terminal domain can facilitate actin binding. By this logic, ERM proteins could be translated in an “ON” state, ready to perform their function.

What pathways localize translationally competent *erm-1* transcripts to the plasma membrane in the early embryo? The pathway that directs (transmembrane protein and secretory protein encoding) mRNAs to the ER is the most well-characterized translation-dependent mRNA trafficking pathway and requires the presence of a signal peptide. As ERM-1 lacks a discernable ER-directing signal peptide and fails to pull down ER associated-components in IP assays, we surmise the pathway that directs translating *erm-1* to the plasma membrane is likely distinct from the ER secretory pathway (Ast et al., 2013; Cohen et al., 2020; Keenan et al., 2001). Alternatively, if *erm-1* is localized in an ER-dependent pathway it would likely be noncanonical.

Future studies will determine the machinery required to traffic *erm-1*, and transcripts like it. A recent study live-imaging *erm-1* mRNA movements reported reduced *erm-1* enrichment at the membrane upon knocking down a dynein motor (DHC-1), suggesting *erm-1* localization could require this component of cytoskeletal trafficking (Li et al., 2021). It will also be interesting to determine whether *erm-1* translation is paused or active during the trafficking process and whether multiple rounds of translation occur at the membrane.

Our screen of FERM and PH-like domain containing genes yielded multiple transcripts with mRNA localization at the plasma membrane suggesting this property is conserved across species. The *ani-1* and *ani-2* ortholog in *Drosophila, anillin*, concentrates at the pseudo-cleavage furrow of Drosophila embryos dependent on translation of both its PH and actin binding domains (Hirashima et al., 2018). (Lee et al., 2020; Parker et al., 2020) Indeed, the expanded use of mRNA imaging and sub-cellularly enriched RNA-seq technologies has led to a greater appreciation that localization of mRNA is a widespread feature of cell biology, not only to the plasma membrane but to a wide diversity of membranes and other subcellular structures (Chouaib et al., 2020; Safieddine et al., 2021). These findings suggest that many proteins may benefit from local translation at their destination site.

## MATERIALS AND METHODS

### Worm husbandry

*C. elegans* strains were cultured according to standard methods (Brenner, 1974). Worm strains were maintained and grown at 20°C on nematode growth medium (NGM: 3 g/L NaCL; 17 g/L agar; 2.5 g/L peptone; 5 mg/L cholesterol; 1mM MgSO_4_; 2.7 g/L KH_2_PO_4_; 0.89 g/L K_2_HPO_4_). Strains used in this study are listed in (**Table S1)**.

### Heat Shock Experiments

Heat shock experiments were performed by transferring harvested embryos to pre-warmed M9 liquid media and incubating at 30°C for 25 minutes. After heat shock, worms were immediately fixed for smFISH.

### RNAi Feeding for smFISH Microscopy

RNAi feeding constructs were obtained from the Ahringer library (Fraser et al., 2000). Bacteria containing inducible dsRNA vectors were grown at 37°C in LB + ampicillin (100 μg/mL) media for 16 hr, spun down and resuspended at 10X original concentration with M9, plated on NGM + Carbenicillin (100 μg/mL) + IPTG (1 mM) plates, and grown at room temperature overnight, or until plates were dry. Embryos were harvested for synchronization from mixed staged worms. Harvested embryos were incubated in M9 for 24 hrs at RT while nutating until all arrived at L1 developmental stage. L1 worms were deposited on RNAi feeding plates and grown at 20°C for 48 hrs. Embryos were harvested from gravid adults and smFISH was conducted. For each gene targeted by RNAi, we performed at least three independent replicates. L4440 RNAi empty vector was used as a negative control and *pop-1* RNAi used as an embryonic lethal positive control. For experiments performing *ifg-1* RNAi, synchronized L1 worms were grown to L2 on OP50 plates before being washed in M9 and transferred to RNAi plates for 48 hrs. *E. coli* strains used in this study can be found in (**Table S2**).

### Permeabilization and drug delivery

For small molecule inhibitor experiments, *perm-1* RNAi was performed to permeabilize the eggshell as described in (Carvalho et al., 2011). Briefly, synchronized L1 worms were fed on RNAi for 48 hrs and embryos were hand-dissected from the treated mothers. Permeabilization by p*erm-1* RNAi was tested by submerging embryos in water to induce bursting from to increased internal osmotic pressure. Additionally, RNAi efficacy was confirmed by *pop-1* RNAi positive control. To harvest embryos, young adult worms were washed off plates in 5ml S-buffer (129 mL 0.05 M K_2_HPO_4_; 871 mL 0.05 M KH_2_PO_4_; 5.85 g NaCl’ 300 +/- 5 mOsm) and allowed to settle to the bottom of a 15 ml conical (no longer than 5min). S buffer was removed, and worms were resuspended in 100 μl S-buffer alone (negative control) or drug diluted in 100 μl S-buffer (500 μg/mL Cycloheximide and 100μg/mL Puromycin final concentrations). The 100 μl drug solution and young adult worms were transferred to a concavity slide, hand dissected in a concavity slide, transferred to a 1.5 mL Eppendorf tube and incubated for 20 min. After incubation with drug solution, 1 mL of -20°C methanol was added to fix the embryos. Embryos were freeze-cracked in liquid nitrogen for 1 min then incubated overnight in methanol at -20°C to continue the fixation. smFISH was performed as described. Due to the fragility of *perm-1* depleted embryos, all spins required in their smFISH preparation were performed at 250 rcf instead of 2000 rcf.

### smFISH

single molecule Fluorescence *In Situ* Hybridization (smFISH) was performed based on the TurboFish protocol (Femino et al., 1998; Nishimura et al., 2015; Parker et al., 2021; Raj and Tyagi, 2010; Raj et al., 2008; Shaffer et al., 2013). Updates specific to *C. elegans* were made using new Biosearch reagents and outlined in (Parker et al., 2021). Using the Stellaris RNA FISH Probe Designer, custom FISH probes were designed for transcripts of interest (Parker et al., 2020; Parker et al., 2021)Biosearch Technologies, Inc., Petaluma, CA) available online at www.biosearchtech.com/stellarisdesigner (version 4.2). Embryos were hybridized with Stellaris RNA FISH Probe sets labeled with CalFluor 610 or Quasar 670 (Biosearch Technologies, Inc.), following the manufacturer’s instructions available online at www.biosearchtech.com/stellarisprotocols. Briefly, adult worms were bleached for embryos, suspended in 1 ml -20°C methanol, freeze cracked in liquid nitrogen for 1 min, and fixed overnight in methanol at -20°C for 1-24 hours. Alternatively, fixation was done at 4 min incubation following the 1 min freeze crack, switching to -20°C acetone incubation for an additional 5 min. ERM-1::GFP worms were freeze-cracked in 1 mL acetone followed by a 35 min incubation at -20°C. After fixation, embryos incubated in Stellaris Wash Buffer A for 5min at RT (Biosearch, SMF-WA1-60) before hybridization in 100 μl Stellaris Hybridization buffer [(Biosearch, SMF-HB1-10) containing 50 pmols of each primer set (up to two channels per experiment) and 10% formamide] 16-48 hrs at 37°C mixing at 400 rpm in a thermomixer (Eppendorf ThermoMixer F1.5). Embryos were washed for 30 min in Stellaris Wash Buffer A, followed by a second wash of Stellaris Wash Buffer A containing 5 μg/ml of DAPI, then 5 min in Wash Buffer B followed by a second 5 min wash (Biosearch, SMF-WB1-20) before incubation in N-propyl gallate antifade (10 mL 100% glycerol, 100 mg N-propyl gallate, 400 μL 1M Tris pH 8.0, 9.6 mL DEPC treated H_2_O) prior to slide preparation. All embryos were centrifuged in spin steps at 2,000 rcf unless otherwise noted. Embryos were mounted using equal volumes hybridized embryos resuspended in N-propyl gallate antifade and Vectashield antifade (Vector Laboratories, H-1000). smFISH image stacks were acquired as described in Parker et al., 2020. on a Photometrics Cool Snap HQ2 camera using a DeltaVision Elite inverted microscope (GE Healthcare), with an Olympus PLAN APO 60X (1.42 NA, PLAPON60XOSC2) objective, an Insight SSI 7-Color Solid State Light Engine, and SoftWorx software (Applied Precision) using 0.2 m z-stacks. Deltavision (SoftWorx) deconvolution software was applied for representative images. Images were further processed using FIJI (Schindelin et al., 2012). For each condition described a minimum of 5 embryos per 4-cell stage, but often many more across multiple cell stages, were imaged. All smFISH and smiFISH probes can be found in (**Table S3**). All raw microscopy images are deposited on Mountain Scholar, a digital, open access data repository associated with Colorado State University Libraries:

### smiFISH

single molecule inexpensive Fluorescence *In Situ* Hybridization (smFISH) was performed as described in (Parker et al., 2021). Briefly, custom primary DNA oligos were designed as described (Tsanov et al., 2016) complementary to the 17 FERM and PH-like domain containing transcripts screened and ordered from IDT (https://www.idtdna.com/pages/products/custom-dna-rna/dna-oligos) (**Table S3**). Secondary FLAPX probes were ordered with dual 5’ and 3’ fluorophore labeling, Cal Fluor 610 or Quasar 670, from Stellaris LGC (Biosearch Technologies, BNS-5082 and FC-1065, respectively). Secondary, fluorophore labeled probes were annealed to primary probes fresh for every experiment in a thermocycler at 85°C for 3 min, 65°C for 3 min, and 25°C for 5 min.

### Quantification of plasma membrane RNA localization

Quantification of transcript localization with reference to the cell membrane was performed as previously described (Parker et al., 2020) using the web application ImJoy (Ouyang et al., 2019). Briefly, RNAs were first detected from raw images using the Matlab code FISH-quant (Mueller et al., 2013). Individual cell outlines were then manually annotated in FIJI for each Z-stack in the micrograph, excluding the uppermost and lowermost stacks where cells are flattened against the slide or coverslip or there is out-of-focus light. The distance of each RNA was then measured from the nearest annotated membrane and binned in 10% distance increments away from nearest membrane to account for change in size between embryos. Total number of RNAs per bin were then normalized by the volume of the concentric areas they occupied. After this normalization, values larger than 1 indicate that for this distance more RNAs are found compared to a randomly distributed sample.

### Quantification of total mRNA

Detection of RNA molecules was performed in the 3D image stacks with FISH-quant (Mueller et al., 2013). Post-processing to calculate the different location metrics was performed as described above with custom written Matlab and Python code. The Python code is implemented in user-friendly plugins for the image processing platform ImJoy (Ouyang et al., 2019). Source code and all scripts used for analysis and figure generation are available here https://github.com/muellerflorian/parker-rna-loc-elegans

To quantify the number of individual mRNAs in the ERM-1[4KN] strain a custom MATLAB script was implemented. FISH-quant detection settings were used to identify candidate mRNA clusters from smFISH micrographs using a Gaussian Mixture Model (GMM). The GMM differentiates independent, single mRNAs from groups of clustered mRNAs by probabilistically fitting a predicted RNA of average intensity and size over each FISH-quant detected RNA.

### Domain search

A smiFISH-based screen was utilized to identify if transcripts with protein domains similar to *erm-1* also displayed membrane localization. WormBase ParaSite was utilized to generate a list of proteins with domain IDs matching those annotated for ERM-1. Proteins with domain IDs matching ERM-1 were subset for genes present in 2-cell stage embryos (Nishimura et al., 2015). This resulted in a list of 149 maternally inherited genes encoding either a FERM or PH-like domain. (**Table S4**).

Candidate genes were further subset using the “Interactive visualizer of differential gene expression in the early *C. elegans* embryo” (http://tintori.bio.unc.edu/, Tintori et al., 2017). Candidate genes were manually curated based on the following criteria: 1) persistence of enrichment in the 4-cell stage embryo, 2) high transcript abundance in the 4-cell stage embryo, 3) homology to genes encoding transcripts with known localization in other biological systems, and 4) existing protein expression data available on Wormbase (https://wormbase.org/) (**Table S5**). Manual curation resulted in 17 candidate genes that are simultaneously maternally inherited and contain FERM or PH-like domains to screen for membrane localization (**Table 1**).

## ACKNOWLEDGEMENTS

We are grateful to Michael Boxem, Susan Mango, Cristina Tocchini, Brooke Montgomery, Tai Montgomery, Jessical Hill, and WormBase for reagents, protocols, equipment, advice, productive discussion, and critical feedback on the manuscript. This work utilized resources from the University of Colorado Boulder Research Computing Group, which is supported by the National Science Foundation (awards ACI-1532235 and ACI-1532236), the University of Colorado Boulder, and Colorado State University. This work utilized microscopy resources from NIH grant 1S10 OD025127 and support from the CSU Microscope Imaging Network. Some strains were provided by the Caenorhabditis Genetics Center, which is funded by National Institutes of Health Office of Research Infrastructure Programs (P40 OD010440). Some figure elements were created in BioRender.

## Funder Grant reference number Author

National Institutes of Health R35GM124877 Erin Osborne Nishimura

NSF MCB CAREER 2143849 Erin Osborne Nishimura

NSF GAUSSI training grant DGE-1450032 Lindsay Winkenbach

NSF GAUSSI training grant DGE-1450032 Dylan Parker

Bridge to Doctorate at Colorado State University 1612513 Robert Williams

## AUTHOR ORCIDS

Lindsay P. Winkenbach https://orcid.org/0000-0002-1766-3260

Dylan M. Parker https://orcid.org/0000-0002-4910-4113

Robert T.P. Williams https://orcid.org/0000-0002-3384-212X

Erin Osborne Nishimura https://orcid.org/0000-0002-4313-4573

## FIGURE LEGENDS

**Table 1: FERM and PH domain-containing, maternally inherited transcripts screened for subcellular localization patterns**. Seventeen transcripts containing either FERM domain or PH-like domains were selected from known maternally inherited mRNAs (Osborne Nishimura et al., 2015, Tintori et al., 2016). Cell enrichment at the 4-cell stage from scRNA-seq data is shown. Transcript localization is briefly described.

